# The chemoreceptor McpA mediates aerotaxis and redox taxis in *Salmonella* Typhimurium

**DOI:** 10.64898/2026.07.03.735341

**Authors:** Kailie Franco, Kendal G. Cooper, Clara Hyunyoung Shin, Olivia Steele-Mortimer, Arden Baylink

## Abstract

Redox environments in nature are shaped by reactive oxygen species (ROS), oxygen, and metal ion chemistry, and exert profound effects on bacterial physiology and survival. In some pathogens, bacterial chemosensory zinc-binding (CZB) proteins mediate responses to inflammatory ROS that may contribute to pathogenesis, but their functions in the many environmental bacteria that also possess them remain unclear. Reports of attraction to hypochlorous acid (HOCl), repulsion from hydroperoxides (ROOH) and other ROS, and responsiveness to exogenous zinc have left unresolved whether CZBs distinguish individual effectors or reflect an underlying sensory logic. Here, using time-resolved video microscopy to directly visualize swimming and taxis behaviors, we show that the CZB-containing receptor McpA in *Salmonella* Typhimurium mediates attraction to O₂ and biologically relevant concentrations of HOCl, H_2_O_2_, and organic hydroperoxides, three types of oxidants generated through the neutrophil oxidative burst. This identifies the CZB domain as a previously unrecognized sensory architecture for aerotaxis, distinct from, and possibly more widespread, than the FAD-dependent Aer and heme-dependent HemAT and Aer2 families. Although Zn²⁺ itself is not redox-active, we show that McpA-dependent taxis is controlled by Zn²⁺ availability and that exogenous O₂ and ROS promote the formation of insoluble zinc precipitates, which may generate Zn²⁺ gradients sensed through the CZB domain. We propose these findings are consistent with a model in which CZBs use soluble, bioavailable Zn²⁺ as a secondary messenger for highly sensitive navigation of environmental redox landscapes.

## Introduction

Approximately 2.4 billion years ago, life on primordial earth was subjected to a mass extinction event driven by the appearance of high levels of molecular oxygen (O_2_), known as the Great Oxygenation Event (GOE) ^1,2^. The increase in atmospheric O_2_ provided opportunities for more efficient metabolism through the use of O_2_ as a terminal electron acceptor, but also new and deadly sources of oxidative stress, such as reactive oxygen species (ROS) ^3^. These chemicals, including the superoxide anion (O_2_^.-^), hydroperoxides (ROOH), and hypochlorous acid (HOCl), can kill cells through lipid peroxidation, DNA damage, base oxidation and deamination, and protein carbonylation ^4–6^. All cells today bear the scars of this evolutionary bottleneck in the form of a sophisticated network of antioxidant enzymes and small-molecule reductants that strictly maintain intracellular redox homeostasis ^7,8^. Cells also engage in redox warfare through the deliberate production of ROS, as exemplified by neutrophils, which generate high concentrations of ROS through the respiratory burst to kill microbial pathogens ^3–6^. The evolution of multiple systems for sensing O₂, ROS, and cellular redox state, including Fnr, ArcAB, SoxR, OxyR, and Aer, reflects the pervasive influence of oxygen availability and oxidative chemistry on cellular function ^7,9–16^.

Chemosensory zinc-binding (CZB) protein domains are found in all major bacterial lineages, suggesting that they originated before the diversification of extant bacterial phyla ^17^. Typically found as components of chemoreceptors and diguanylate cyclases, CZB domains serve as regulators of chemotaxis and c-di-GMP signaling, thereby controlling behaviors central to motility and navigation, lifestyle transitions, and colonization ^17,18^. As their name suggests, CZB domains bind a single Zn^2+^ ion with femtomolar-range affinity, ligated by a 3His, 1Cys motif that was first identified as a conserved domain of cytoplasmic chemoreceptors ^19^, and characterized structurally and biochemically through studies of the *Helicobacter pylori* chemoreceptor TlpD and the *Escherichia coli* diguanylate cyclase DgcZ ^17,20–23^, yet the biological purpose and significance of their zinc-sensing function has remained enigmatic ^17–20,22–24^. Although femtomolar Zn²⁺ sensitivity has been reported for bacterial metalloregulators such as Zur and ZntR, cytosolic Zn²⁺ is buffered by competing protein and small-molecule ligands whose relative affinities and exchange kinetics determine the labile, bioavailable pool; this dynamic zinc speciation is difficult to recapitulate *in vitro*, making it challenging to connect CZB zinc binding to physiological responses ^25–27^. Prior studies have demonstrated that CZB proteins mediate behavioral responses to ROS, and the Zn^2+^ is exchangeable, but seemingly contradictory results have made the biological function difficult to resolve ^17,22–24,28,29^. TlpD arguably stands as the best-characterized CZB protein to date, and confers strong attraction to HOCl *in vitro*, paradoxically navigating the bacteria toward a potent bactericide ^21,22,30^. Whether this behavior is artifactual and/or a species-specific phenomenon is unknown. Further, some studies have proposed TlpD mediates repulsion from H_2_O_2_ and other inflammatory ROS ^24,28^. Whether CZB proteins are imbued with chemoselectivity that orchestrates discrete behaviors to these different stimuli, or if there exists an underlying coherent sensory logic linking these responses, has so far remained unresolved.

A current mechanistic model of CZB sensory function posits that the Cys-Zn moiety is sensitive to direct oxidation by neutrophilic HOCl to form cysteine-sulfenic acid (Cys-SOH), resulting in detachment of the Cys from the zinc-binding core and a local unfolding of the domain that promotes release of the Zn^2+^ ion and signal transduction ^17,22^. Interestingly, at the protein level, the conserved zinc-binding Cys does exhibit chemoselectivity and is preferentially oxidized by HOCl and inefficiently oxidized by hydroperoxides ^17,22,23^. This mechanism of direct cysteine oxidation proposed for CZBs is analogous to that seen for zinc-finger proteins and other systems shown to undergo redox-dependent zinc release ^17,22,23,31^, but some lines of evidence argue against this model reflecting a universally conserved function of CZBs. First, the origin of CZBs likely predates animals and they are possessed by diverse bacteria, including environmental species not associated with hosts that would rarely encounter concentrated hypohalous acids like those that occur at sites of inflammation ^17^. Second, we recently showed that while the zinc-binding Cys mediates bacterial fitness *in vivo* during colitis, a disease state where the enteric environment has increased oxygenation as well as prevalent HOCl and other host-derived ROS, infection in a mouse model deficient in HOCl generation did not impact CZB-mediated colonization ^32–35^.

A major obstacle to defining the function and sensory mechanism of CZB domains is the lack of a clear understanding of the behavioral outputs they direct. Most studies have been conducted in *H. pylori*, which poses challenges for dissecting chemotaxis mechanisms due to its variable and fickle motility, limited genetic tractability, and requirement for growth in rich liquid media. These challenges limit the ability to perform chemotaxis assays in a defined chemical background and to clearly identify the effector species being sensed ^22,24,29,36,37^. Moreover, redox-active effectors are inherently difficult to study because they are transient and experiments require careful control of the atmospheric background, which is technically challenging and not typically undertaken.

Herein, we employed a more genetically tractable and experimentally accessible model system that enables live-imaging of behavioral responses to test different models of CZB sensory mechanisms. We investigated the CZB-regulated chemoreceptor methyl-accepting chemotaxis protein A (McpA) from *Salmonella* Typhimurium, an orthologue of TlpD with similar domain architecture and approximately 20% sequence identity ^18,22,32^. We employed a chemotaxis platform that enables direct visualization of bacterial redox taxis in response to injected effectors under controlled atmospheric conditions, and, orthogonally, a high-throughput swimming assay with automated tracking and analysis of hundreds to thousands of individual bacteria. Using these complementary approaches, we propose a model of redox sensing mediated through depletion of soluble Zn^2+^ that may represent the conserved and ancestral function of CZB proteins.

## Results

### Salmonella Typhimurium exhibits concentration-dependent taxis to ROS

During infection, *Salmonella* Typhimurium induces colitis and leverages metabolic advantages to thrive in the inflamed gut, an environment where infiltrating neutrophils generate large quantities of ROS, transforming the redox environment ^33,34,38–40^. Through NADPH oxidase, neutrophils generate H₂O₂ that is subsequently converted to the strong oxidant HOCl by myeloperoxidase; activated neutrophils can generate HOCl at rates approaching 134 mM min⁻¹, and local HOCl concentrations within inflamed tissues or phagosomes have been estimated to reach up to 5 mM ^4,41^. The oxidative burst also generates organic hydroperoxides (ROOH) via lipid peroxidation, highly reactive oxidants capable of inducing cell lysis ^42,43^. Hence, *Salmonella* regularly experiences high concentrations of these bactericidal oxidants, each possessing different reactivities, half-lives, and preferred oxidation targets, and no prior studies have tested if McpA mediates taxis to these oxidants.

To address this, we employed the Chemosensory Injection Rig Assay (CIRA, Fig. 1A), which uses a glass microcapillary to continuously inject femtoliter volumes of effectors into a suspension of motile bacteria and monitor taxis responses through video microscopy ^22,44–46^. In these experiments, swimming bacteria initially have a homogenous distribution, and exposure to treatment results in either no discernible change (null response), an influx of bacteria toward the treatment source (chemoattraction), or decrease in bacteria (chemorepulsion). Modeling of ROS diffusion in this assay predicts a steep, orders-of-magnitude concentration gradient from the source, such that even a 1 mM injection exposes most bacteria within the field of view to physiologically relevant pico-to nanomolar ROS concentrations (Fig. 1B).

**Fig. 1.**
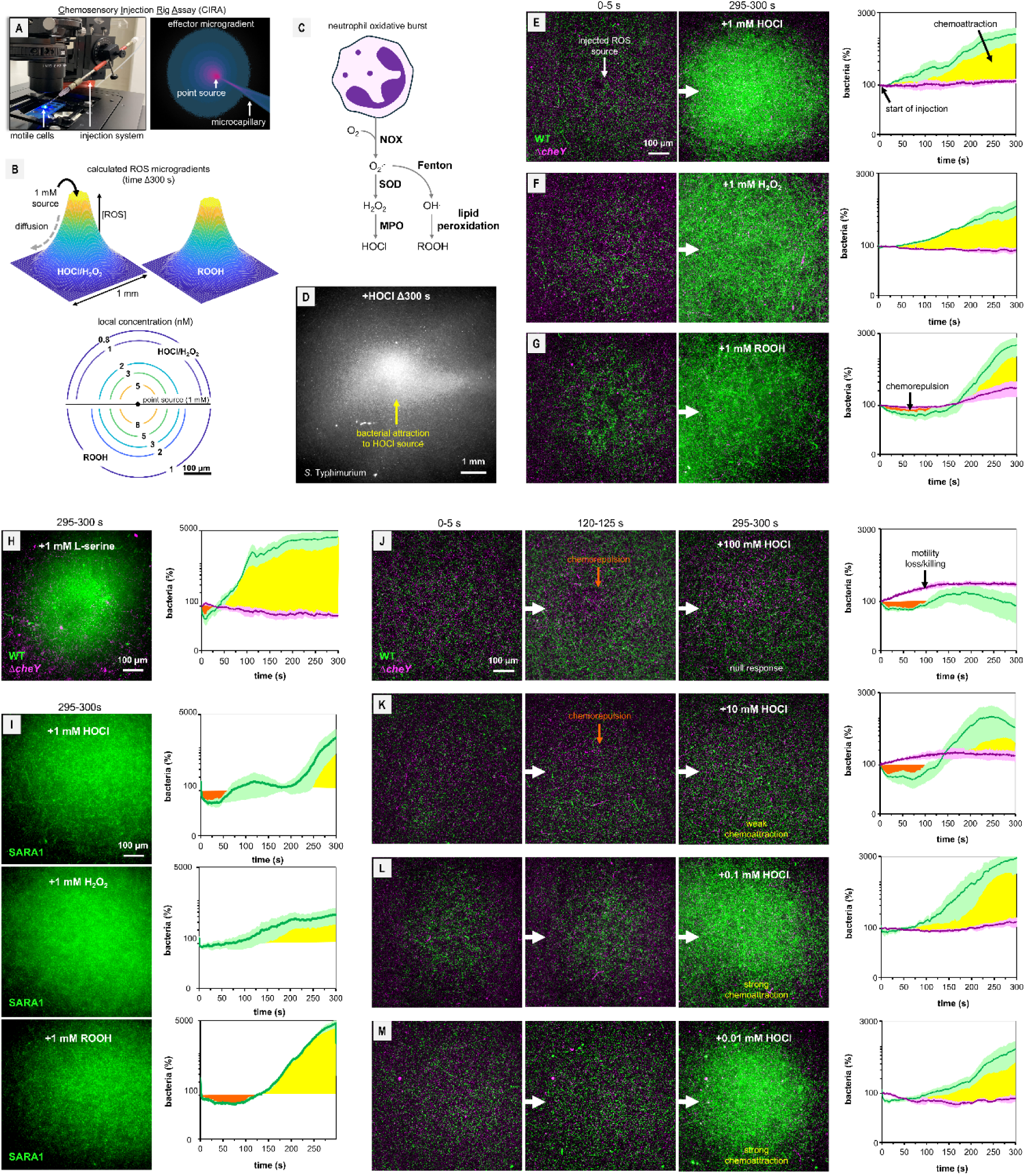
Neutrophil-derived reactive oxygen species drive chemoattraction in S. Typhimurium. A. Experimental design of CIRA. B. Calculated diffusion models for the local concentrations of reactive oxygen species (ROS) experienced by bacteria at a given distance from the injection source at time 300 s. Flow of the injection is continuous at approximately 300 femtoliters per minute. C. Overview of ROS production by neutrophils through the oxidative burst. D. Widefield view of S. Typhimurium IR715 attracted to a 1 mM HOCl injected treatment using CIRA at time 300 s. E-G. Competition experiments with WT *S.* Typhimurium IR715 (green) and an isogenic Δc*heY* mutant (magenta), responding to injected ROS treatments as indicated. Representative max projections over 5 s are shown at the experiment start (left) and end (center). Quantification of relative cells in the field of view are shown right. Chemoattraction responses are highlighted in yellow throughout, and chemorepulsion responses in dark orange, defined as the number of cells in view exceeding or below the initial starting levels, respectively, and not due to motility loss or killing. Plots show the relative number of cells in the field of view as a percentage of the number at the start of treatment (time 0 = 100%). H. Chemoattraction of IR715 to L-Ser. I. Chemoattraction of S. Typhimurium strain SARA1 to neutrophil ROS. J-K. Taxis of IR715 to different source concentrations of HOCl. Graphs depict means with error bars depicting standard error of the mean (SEM) of 4-5 replicate experiments.

We employed CIRA to model the neutrophil respiratory burst and study taxis responses to discrete ROS (Fig. 1C). We performed competition assays with fluorescently-tagged WT *Salmonella* Typhimurium IR715 (IR715 WT) and a *cheY* knockout mutant, which is motile but blind to chemoeffector stimuli, and viewed their taxis responses simultaneously (Table S1). CIRA experiments using 1 mM sources of HOCl, H_2_O_2_, and cumene hydroperoxide, which we used as a model organic hydroperoxide, all elicited chemoattraction, with more WT cells coming into the field of view over time, albeit a more diffuse response than seen for L-Ser, a well-characterized chemoattractant (Fig. 1D-H, Video S1) ^44,45^. As *S. enterica* strains can exhibit rather different phenotypes, even among the same serovar, we also tested ROS taxis in *S.* Typhimurium strain SARA1, and also found this clinical isolate to be attracted to all three oxidants (Fig. 1I). Importantly, the distribution of the *cheY* cells are not impacted by ROS treatments, and so the behavior cannot be due to a trapping mechanism or impairment of motility leading to aggregation of cells near the ROS source. These findings revealed that, like seen previously for *H. pylori*, *S.* Typhimurium is attracted to HOCl and also uncovered a previously unknown attraction to hydroperoxides ^22^.

While it may seem counterintuitive that bacteria are attracted to sources of bactericidal oxidants, the consistency of this response across all three neutrophilic ROS led us to ask what concentrations elicit taxis. We performed CIRA experiments across a wide range of physiologically relevant HOCl concentrations and found responses to be concentration-dependent and adaptive (Fig. 1J-M). At a source concentration of 100 mM, WT cells are initially repelled and then adapt after about four minutes and return to a random distribution (Fig. 1J). At 10 mM this response is similar with a shorter adaptive period, with WT cells exhibiting chemorepulsion until approximately 2.5 minutes, and, thereafter, chemoattraction (Fig. 1K). HOCl sources of 0.01-1 mM all promoted clear chemoattraction (Fig. 1E, L-M). Hence, *S.* Typhimurium mediates differential taxis based on the concentration of exogenous ROS, with chemoattraction occurring only to levels in the nanomolar to picomolar range.

### McpA and other chemoreceptors regulate swimming behavior in response to HOCl

Having established that *S.* Typhimurium is attracted to ROS sources through chemotaxis, we next sought to identify chemoreceptors responsible for regulating swimming behavior in response to exogenous HOCl. In the canonical model of chemotaxis, swimming bacteria alternate between runs and tumbles, and chemoattractants elicit decreased tumbling rates, which is also reflected in increased migration via longer uninterrupted runs (Fig. 2A) ^30,47–52^. To assess these behaviors in response to HOCl, we employed an assay in which bacteria are bathed in a homogenous treatment and then videos of swimming bacteria are collected and processed through an automated tracking pipeline to plot hundreds-to-thousands of individual bacterial swimming trajectories and quantify migration, speed, and tumbling frequency (Fig. 2A) ^22,53^. We performed this assay with WT and single chemoreceptor mutants to identify those involved in mediating HOCl taxis.

**Fig. 2.**
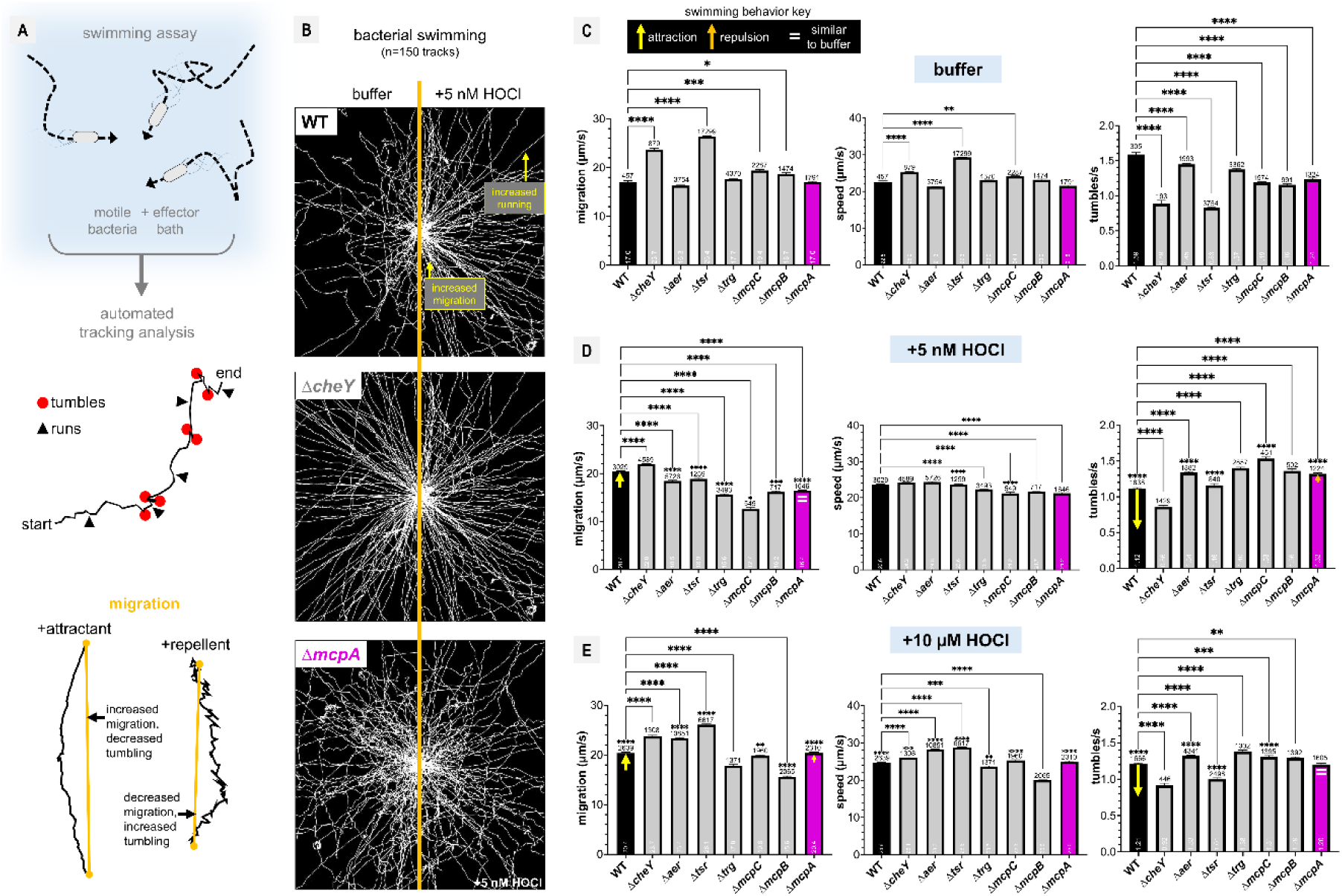
Bacterial swimming behavior is responsive to neutrophil ROS. A. Overview of swimming assay in which bacteria (*S.* Typhimurium IR715) are immersed in a bath of chemoeffector, and swimming trajectories are recorded for 30 s by video microscopy. Tracks are analyzed for tumble rate and migration as metrics of attraction or repulsion behavior, with swimming speed serving as a control for impacts on motility. B. Representative plots of bacterial swimming trajectories are shown (n=150) in buffer (left panels) or with addition of 5 nM HOCl (right panels). When exposed to HOCl, the WT tracks are visibly smoother, reflecting increased migration and decreased tumbling, whereas Δ*mcpA* is unchanged. C-E. Quantifications of bacterial migration (left), swimming speed (center), and tumbling rate (right) under different treatment conditions, as indicated. For clarity, key behavioral changes of the WT and Δ*mcpA* upon addition of effector corresponding to taxis behaviors are noted with either yellow arrows (attraction behavior), orange arrow (repulsion behavior), or = to indicate little or no change, in relation to that strain’s behavior in buffer; the length of the arrows corresponds to the magnitude of change. Graphs depict means with error bars as standard error of the mean (SEM) of 5-8 replicate experiments; numbers above bars denote tracks, while numbers within the bar are population means. Statistical significances between strains, denoted by bars within each treatment group, were calculated using a Kruskal-Wallis test followed by Dunn’s multiple comparison tests, compared to the WT strain within that treatment group. Statistical significances for strain differences compared to buffer treatment version of the same strain are denoted by stars above the number of tracks and were calculated by unpaired t-tests (not significant, not noted; * p < 0.05, ** p < 0.01, *** p < 0.001, **** p < 0.0001).

In these experiments, we saw that WT swimming tracks appear more similar to Δ*cheY* upon treatment with 5 nM HOCl, with fewer direction changes, indicative of increased running, while Δ*cheY* and Δ*mcpA* have similar tracks regardless of treatment (Fig. 2B). WT cells in the presence of 5 nM and 10 µM HOCl, concentrations in the range we calculate to stimulate chemoattraction in CIRA, shows increased migration and decreased tumbling rate, consistent with chemoattraction (Fig. 2B-E). Swimming speed was similar across treatments, indicating HOCl at these concentrations does not impair motility (Fig. 2C-E). We also found that many receptor mutants show some degree of defect in overall taxis and also in HOCl response. *mcpA, aer,* and *trg* lost HOCl-responsiveness, as migration and tumbling rate is similar across treatments, while *tsr, mcpB,* and *mcpC* show elevated tumble rates, i.e. an inverted response (Fig. 2C-E). These defects in HOCl taxis in these mutants could be either from loss of direct HOCl-sensing or perhaps altered function of the chemoreceptor nanoarray; Aer and Tsr have also previously been implicated in ROS taxis and aerotaxis ^33,54–56^. Together, in terms of regulation of swimming behavior in response to HOCl exposure, we see an attraction phenotype at low concentrations that is consistent with the localization toward ROS sources in CIRA, and multiple receptors, including McpA, contribute to this behavior, although indirect effects of receptor deletion cannot be excluded.

### Salmonella Typhimurium tolerates ROS concentrations that elicit chemoattraction

The differential chemotactic responses across ROS concentration regimes led us to investigate how these ROS concentrations impact the growth of *S*. Typhimurium. Bacterial growth was assessed under both normoxic conditions, where aerobic respiration predominates, and hypoxia (1% O₂ + 10% CO₂), mimicking the low-oxygen environment of the intestinal lumen ^57,58^. As expected, higher O_2_ levels increase bacterial growth of this facultative anaerobe (Fig. 3A). Of the neutrophil-derived ROS, HOCl is the most potent bactericide, with IC_50_ values of 8.9 µM and 36.6 µM under hypoxic and normoxic conditions, respectively (Fig. 3A-B). Hydroperoxides are better tolerated with high micromolar to millimolar IC_50_ values (Fig. 3A-B). Interestingly, under hypoxic conditions, low levels of H_2_O_2_ conferred a small growth benefit during the stationary/death phases of growth (Fig. 3B). This observation is consistent with prior work showing *S.* Typhimurium can utilize H_2_O_2_ as an alternative electron acceptor under oxygen-limiting conditions ^34,35,59,60^. Together, these data reveal that *S.* Typhimurium is attracted to low ROS levels it can tolerate, or even obtain a growth advantage from, and is repelled at high ROS concentrations that are inhibitory to growth.

**Fig. 3.**
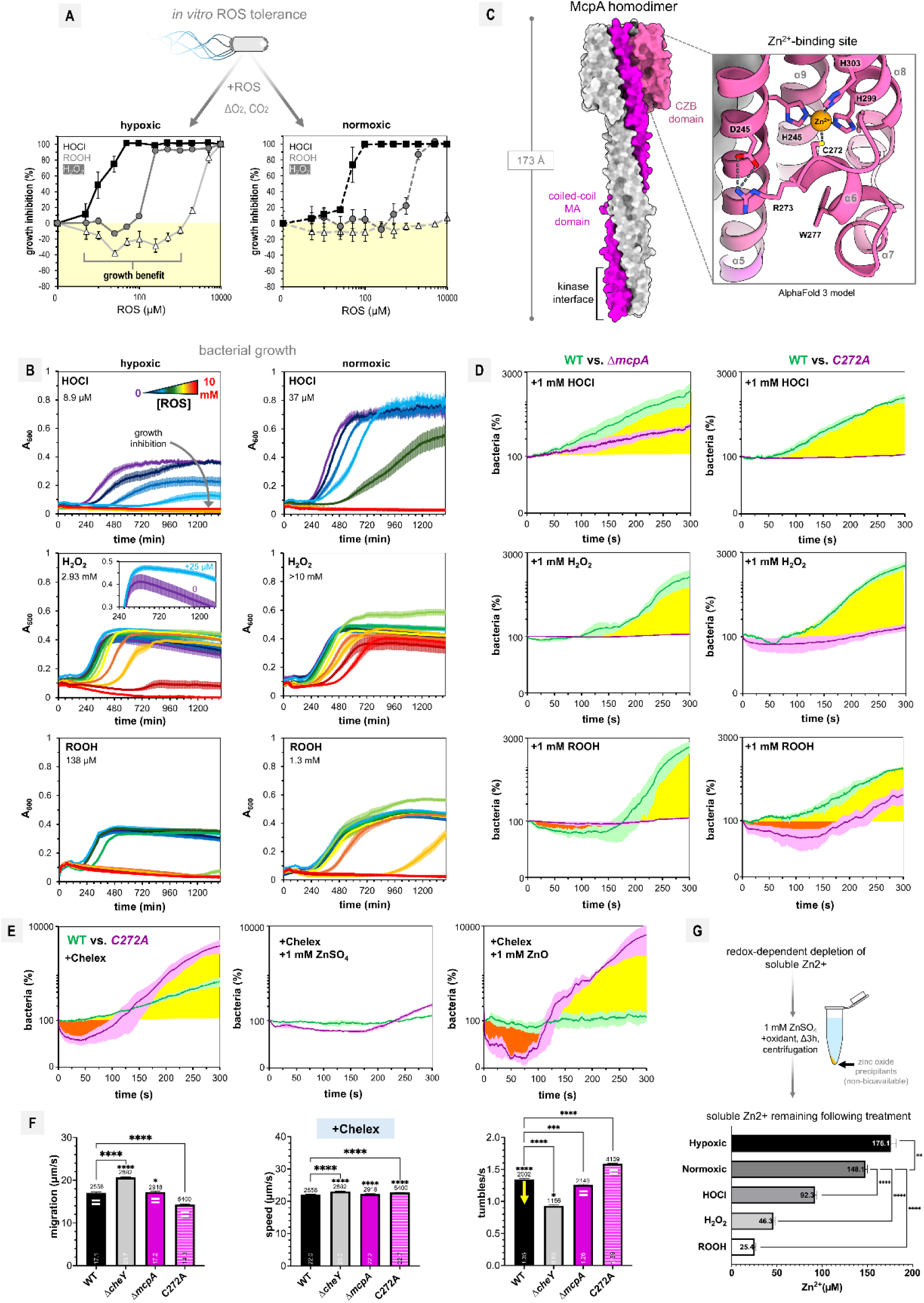
McpA mediates taxis to ROS and Zn^2+^ gradients A. Shown is the relative growth inhibition of *S.* Typhimurium IR715 growth inhibition in the presence of different ROS under either normoxic (atmospheric) or hypoxic (1% O_2_ + 10% CO_2_) conditions at 37 °C (n=24). Growth enhancement is highlighted in pale yellow. Relative growth was calculated based on the integrated area of the curve for each set of growth curve data relative to the buffer-treated experiments. B. Growth curves for ROS titrations shown in A, with calculated half-maximal inhibitory values (IC_50_) noted in top left corner of plots; H_2_O_2_ hypoxic plot inset shows growth enhancement in presence of 25 µM H_2_O_2_ (blue) over buffer (purple). Increasing ROS addition is depicted on a rainbow color scale from 0 (purple) to red (10 mM). C. Alphafold3 model of the McpA homodimer with molecular surface rendering. One monomer is shown in white, and the with its coiled-coil MA domain in magenta and CZB domain in pink. The inset shows a closeup view of the predicted zinc-binding site, with key residues denoted. Secondary structure assignments are based on the crystal structure of TlpD from *H. pylori* ^18^. Dashed gray lines indicate predicted hydrogen or zinc-ligation interactions. For clarity, an N-terminal region of about 40 residues that is predicted to be unstructured is not shown. D. CIRA competition experiments with WT and Δ*mcpA* (left) or Δ*mcpA* pmcpA C272A (right, denoted as *C272A* for clarity) responding to injected ROS, as indicated. WT is green and mutants are magenta. E. Competition CIRA with WT (green) and C272A (magenta) in response to either an injected source of Chelex-treated buffer (left), or with Chelex-treated buffer supplemented with 1 mM ZnSO_4_ (center) or 1 mM ZnO (right). F. Swimming behavior in the presence of Chelex-treated buffer, which substantially lowers to free Zn^2+^ ion concentration. G. Shown top is an experimental overview for quantification of redox-mediated depletion of soluble free Zn^2+^ in solution. Note that these are not the concentrations present in CIRA and swimming experiments but instead reflect the bioavailable Zn^2+^ ion remaining in a 1 mM solution of ZnSO_4_ after oxidant treatment and centrifugation to remove non-bioavailable precipitants. Unless otherwise indicated, data means, error bars, n, and statistical calculations are calculated as in Fig. 1-2.

### McpA and its zinc-binding Cys are required for ROS chemoattraction

As McpA was one of the receptors confirmed to be required for regulation of swimming behavior in response to HOCl, we next wanted to determine if this receptor is required for attraction to HOCl sources and other neutrophil-derived ROS (Fig. 3C). Using CIRA, we performed competition experiments with WT and Δ*mcpA* to 1 mM sources of HOCl and hydroperoxides, and found McpA to be necessary for full chemoattraction to HOCl, and hydroperoxide chemoattraction was completely dependent upon McpA (Fig. 3D, Video S2). As a control, we also tested HOCl taxis for a genetic complement of McpA, with the *mcpA* gene integrated at the glmS locus under its own promoter ^32^, and confirmed a similar phenotype as seen for WT, although there was only a partial recovery of activity (Fig. S1A).

To address whether ROS-sensing is mediated through the CZB zinc-binding site we employed a chromosomally-inserted C272A mutant ^32^, the cysteine residue universally conserved across all CZB proteins that coordinates the Zn^2+^ ligand and the target of direct HOCl oxidation in other systems (Fig. 3C) ^17,22,23^. In prior *in vitro* work, substitution of the conserved Cys was shown to reduce Zn²⁺-binding affinity by approximately an order of magnitude; however, because the binding affinity remains in the femtomolar range, the Zn^2+^ ligand is retained in the absence of a strong competing chelator ^17,20^. Using CIRA, we found that the C272A mutant lacks responsiveness to HOCl and shows diminished attraction to hydroperoxides (Fig. 3D, Video S3). Interestingly, the C272A mutant actually shows a stronger defect in HOCl chemoattraction than Δ*mcpA*, but the reason behind this is unclear (Fig. 3D). Further studies of swimming behavior confirmed the C272A mutant to have defects in responsiveness to ROS in terms of migration and tumble rate, and some assays also showed phenotypic differences between the Δ*mcpA* and C272A mutant (Fig. S1B-D). For example, H_2_O_2_ reduces tumbling rate for both WT and Δ*mcpA* but increases tumbling in the C272A mutant (Fig. S1C). Hence, while the C272A mutant was found to have impaired ROS taxis, it was not a phenocopy of the Δ*mcpA*, which may be explained by the cysteine substitution weakening the Zn^2+^ affinity of the binding site rather than rendering it completely non-functional ^20^. Together, these data identify McpA as a mediator of ROS chemoattraction in *S.* Typhimurium and implicate the conserved zinc-binding Cys in ROS taxis.

### Elevated O₂ and ROS decrease the concentration of soluble Zn²⁺

The hypothesis that CZBs act as HOCl-sensors through preferential direct oxidation of the zinc-binding Cys thiolate predicts McpA would mediate taxis only to HOCl and not hydroperoxides that have are poorly reactive with the Cys-Zn moiety ^17,22,23,30^. That we observe similar behavioral responses to both HOCl and hydroperoxides dependent upon C272 argues there must exist a different mechanism that mediates broad sensing of ROS. This led us to consider whether the sensory function of McpA might involve redox-dependent changes in exogenous Zn²⁺ availability.

To test this hypothesis, we first assessed whether McpA is responsive to changes in exogenous Zn^2+^ ion levels; we expected this to be the case given that exchange of the CZB-bound Zn²⁺ ion has been demonstrated *in vitro* and DgcZ-mediated biofilm formation responds to changes in exogenous Zn²⁺ levels ^17,20^. Because Zn^2+^ is essential for cell viability, measurement of chemotaxis in fully Zn^2+^-depleted conditions is not possible, as cells rapidly lose motility (data not shown). However, we found that addition of tolerable, low micromolar levels of the divalent metal ion-chelating agent EDTA to the chemotaxis buffer nullifies attraction to HOCl, and sometimes results in repulsion close to the oxidant source (Fig. S1E). To investigate how cells respond to gradients of Zn^2+^, we used CIRA and assessed taxis to Chelex-treated buffer, which substantially depletes divalent metal ions such as Zn^2+^ ^61^. In these experiments, we again saw the WT exhibit chemoattraction, and notably, that the C272A mutant exhibits an even greater attraction to zinc-depleted buffer (Fig. 3E, Video S4). One possible explanation for this behavior is that the introduction of a local exogenous depletion in Zn^2+^ concentration drives the intracellular zinc equilibrium to promote release of Zn^2+^ by McpA, mediating an attraction response, and that the weaker Zn^2+^ binding affinity of the C272A mutant results in a stronger effect.

When Zn^2+^ was supplemented back into Chelex-treated buffer in the form of ZnSO_4_, WT and C272A were no longer responsive to the treatment (i.e., a null rather than a repellent response) (Fig. 3E, Video S4). We also tested supplementation with ZnO and found in that case WT lost attraction, but C272A still demonstrated attraction to the buffer (Fig. 3E, Video S4). One possible explanation is that ZnO generates a small pool of soluble Zn^2+^ sufficient to restore Zn^2+^ binding by the high-affinity WT receptor and abolish attraction, whereas these concentrations remain below the sensory threshold of the lower-affinity C272A mutant, allowing it to retain attraction to the treatment. In terms of swimming behavior, zinc depletion slightly lowers tumbling rate dependent upon Δ*mcpA* and C272A, consistent with an attraction response to low Zn^2+^ (Fig. 3F). These findings together are consistent with McpA mediating responses to alterations in exogenous Zn^2+^ sans any ROS addition, and also suggest that the sensory system may discriminate between free Zn^2+^ (aq.) and other zinc species.

Although Zn²⁺ itself is not redox active, environmental redox conditions influence zinc speciation, or the distribution of zinc between free, aqueous Zn^2+^ and insoluble or complexed forms, by shifting the equilibrium between soluble Zn²⁺ and insoluble zinc-containing precipitates ^62–66^. Given the responsiveness of McpA to exogenous Zn^2+^ concentrations, we hypothesized that Zn²⁺ could function as a secondary messenger for environmental redox potential, with oxidizing conditions depleting the extracellular soluble Zn^2+^ pool through precipitation to insoluble zinc species that are either not recognized by McpA and/or are not bioavailable. To test if these oxidants impact Zn^2+^ ion availability in solution, we prepared aqueous solutions of 1 mM ZnSO_4_ and treated them with HOCl, H_2_O_2_, or cumene hydroperoxide, and compared solutions incubated at normoxic or hypoxic conditions to test the influence of atmospheric O_2_ levels (Fig. 3G). After three hours, solutions were centrifuged to remove zinc precipitants and the remaining soluble Zn^2+^ ion content in solution was determined using the Zn^2+^-specific probe Zinpyr-1 ^17,67^.

These experiments revealed that hypoxic solutions retain the highest amount of soluble Zn^2+^, followed by normoxic buffer, and ROS addition reduces the soluble Zn²⁺ pool even further by promoting formation of insoluble zinc species (Fig. 3G). Although the precise composition of these precipitates was not determined, they may include zinc oxides, zinc hydroxide, zinc phosphates, and zinc oxyhydroxide species ^68,69^. Thus, there appears to be a reciprocal relationship with oxidant addition and depletion of soluble Zn^2+^ ion (Fig. 3G).

### McpA mediates positive aerotaxis under hypoxic conditions

After learning that McpA is responsive to exogenous Zn^2+^ ion levels in solution, and that this can be modulated not only by ROS but also by O_2_, we wondered if McpA might mediate taxis in response to different levels of oxygenation, i.e. aerotaxis ^70–72^. Chemotaxis experiments are typically performed under normoxic (i.e., 21% O_2_) conditions, which obscures aerotaxis behaviors, and regulating oxygen content can be challenging or incompatible with certain assays. To overcome this limitation, we built a custom hypoxic chamber to house our CIRA setup, enabling real-time visualization of aerotaxis in response to microinjected sources of defined effectors (Fig. 4A-C).

**Fig. 4:**
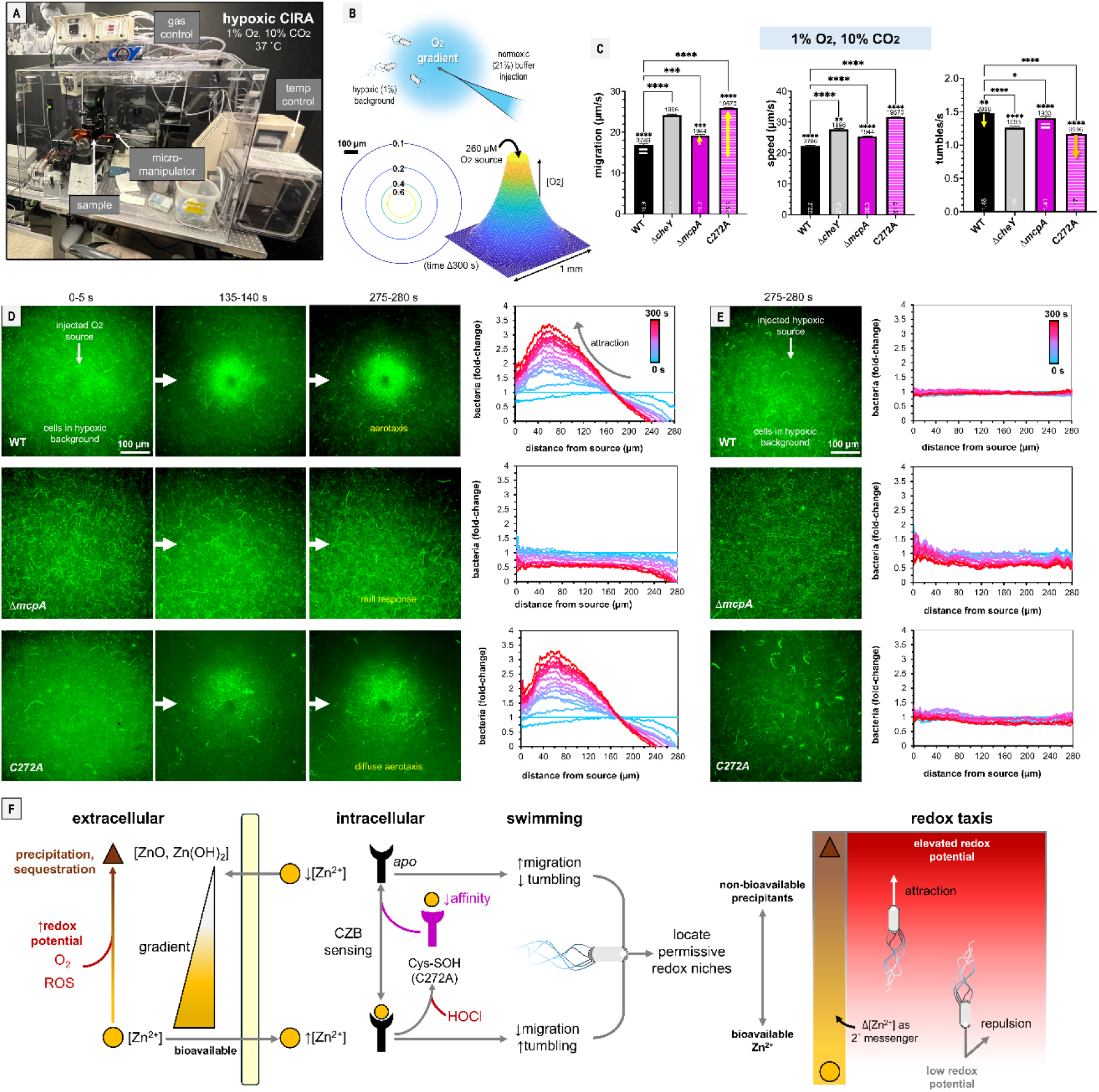
McpA mediates aerotaxis. A. Experimental setup for hypoxic CIRA. The injection apparatus is housed within a hypoxic chamber for atmospheric and temperature control. Cell cultures are grown overnight under the desired conditions prior to experimentation. For clarity, the chamber is shown with its front panel removed. B. Shown top is the experimental schematic to test aerotaxis, in which motile cells under hypoxic conditions that mimic the intestinal environment (1% O_2_, 10% CO_2_, 37 °C) in which motile cells under hypoxic conditions are exposed to injection of normoxic buffer solution. Shown below is the calculated O_2_ diffusion gradient at time 300 s, assuming a 260 µM O₂ source, corresponding to the dissolved O_2_ concentration expected at 21% atmospheric O_2_ based on Henry’s law. C. Bacterial swimming under hypoxic conditions, with behavioral indicators (arrows, =) relative to swimming in normoxic buffer. D. CIRA showing taxis for cells under hypoxic conditions in response to injected normoxic buffer. The plots shown on the right quantify the relative distribution of the bacterial population in relation to the treatment source over time. The initial unbiased population distribution in these plots is indicated with a blue line (time 0), with the mean distributions between at 20 s internals shown as a blue-to-red spectrum at 20 s intervals, with the experiment end at 300 s. E. Experiments as shown in D but with injection of hypoxic-equilibrated buffer, showing a null response. F. Model of CZB sensory function and use of redox-mediated Zn^2+^ speciation as a secondary messenger for redox taxis. Oxidants promote the formation of insoluble Zn^2+^ precipitants (brown triangle), depleting the local Zn^2+^ pool and establishing a concentration gradient of soluble Zn^2+^ ion (orange circle). The change in the extracellular zinc equilibria perturbs the intracellular Zn^2+^ pool, which is sensed through the CZB domain through direct binding of Zn^2+^. When bound to Zn^2+^, McpA promotes increased tumbling and decreased swimming migration, facilitating chemorepulsion. When Zn^2+^ is low, the McpA population shifts toward a zinc-free *apo* state, which decreases tumbling and increases swimming migration, facilitating chemoattraction. For bacteria exposed to high concentrations of HOCl, there is also the direct oxidation mechanism in which the zinc-binding Cys is reversibly oxidized to a cysteine sulfenate (Cys-SOH), which lowers affinity for Zn^2+^ and promotes zinc release, and is modeled by the C272A mutant (magenta). At the cellular and population levels, this enables *S.* Typhimurium to perform redox taxis, whereby McpA directs attraction to regions with low bioavailable Zn^2+^, coinciding with sources of O_2_, and certain types of ROS, that can provide metabolic or other advantages. Data, error bars, and statistical tests are as described above.

We used this setup to test the response of cells cultured under hypoxic conditions that approximate the low-oxygen conditions of the gut environment of *Salmonella* (1% O_2_ and 10% CO_2_) to an injected source of normoxic buffer (Fig. 4B). *S.* Typhimurium exhibits strong attraction to elevated O_2_ that is dependent upon McpA, with the C272A mutant retaining attraction similar in overall magnitude to the WT but with a more diffuse cell population (Fig. 4D, Video S5). In contrast, cells exhibit no response to injected hypoxic buffer, supporting that the attraction is a response to the elevated O_2_, and presumably the locally lowered Zn^2+^ ion concentration (Fig. 4E). Swimming behavior is relatively consistent with the results from CIRA (Fig. 4C). Under hypoxic conditions, the WT shows lower migration and higher tumbling rate than under normoxic conditions indicative of a repulsion from lower O_2_ levels; *mcpA* has slightly higher migration and lower tumbling rate than WT, supporting a role for McpA in mediating the swimming behavior, and C272A had migration and tumbling rate similar to *cheY*, suggesting, potentially, that its lowered zinc-binding affinity is less able to distinguish the differential Zn^2+^ levels in hypoxic versus normoxic conditions, although tumbling rate is indeed lowered for this mutant under hypoxia (Fig. 4C). Together, these results suggest that McpA mediates aerotaxis toward elevated O_2_. As expected, normoxic conditions provide a substantial growth benefit for the bacterium, providing a biological rationale for this behavior (Fig. 3A-B).

Based on these data, we propose a working model in which McpA senses Zn^2+^ as a secondary messenger of environmental redox state (Fig. 4F). Oxidizing conditions driven by O₂ or ROS shift zinc speciation toward insoluble, biologically unavailable zinc precipitates, thereby depleting the soluble Zn²⁺ pool sensed by McpA, promoting an *apo* receptor state and chemoattraction. This ultimately serves to drive attraction of the bacteria toward environments with appropriate redox potential permissive for survival (Fig. 4F). As the structural elements required for this sensing mechanism are conserved across all CZB orthologs, and because the ability to identify favorable redox environments is presumably advantageous across diverse ecological niches, it is possible that redox taxis is the conserved and ancestral function of CZB-domain signaling proteins.

## Discussion

At an early period of evolution, bacteria were confronted with a dramatic elevation in oxygenation, with both enriched O_2_ and ROS levels imposing new selective pressures through their propensity to induce oxidative damage ^1–3,6^. CZB proteins provide an interesting window into how primordial bacteria began to sense the local redox landscape and seek oxidants for their metabolic benefit ^7,8^. Because CZB proteins are hypothesized to have originated before the GOE, the subsequent rise in atmospheric O_2_ may have conferred a selective advantage on this mode of redox sensing, contributing to the widespread conservation of CZB proteins across bacterial phyla ^17,18^. This work establishes that McpA, a CZB-regulated chemoreceptor, mediates aerotaxis, identifying the CZB domain as a previously unrecognized sensory architecture for O₂ taxis. Although the FAD-dependent Aer and heme-dependent HemAT families have been characterized as O₂ sensors and/or aerotaxis receptors in diverse bacteria, their phylogenetic distributions appear to be more restricted than that of CZB proteins: Aer is concentrated primarily in Proteobacteria and Spirochaetota, while HemAT is found predominantly in Bacilli and Clostridia ^73^. By comparison, TlpD-like CZB chemoreceptors have been identified in 1,223 bacterial species, and CZB domains occur across 21 bacterial phyla and six candidate phyla ^17,21^. The broad phylogenetic distribution of CZB proteins, together with the discovery that McpA mediates aerotaxis, suggests that CZB sensory systems may represent one of the most widespread O₂-sensing architectures among prokaryotes.

Cells exert tight control over cytosolic redox reactions and survive within a relatively narrow regime of tolerated redox stress, which differs among cell types and species that inhabit different environments ^6–8,10^. Thus, it is conceivably of general importance for bacteria to possess means to sense local redox potential and identify niches suitable for their particular metabolism. In terms of oxidants mediating metabolic advantage, an especially pertinent example is O_2_, but there is an emerging understanding that at low levels certain ROS can also serve as electron donors and promote metabolism; others may serve as cues that do not themselves impart growth advantages but signal favorable environments, such as damaged tissue, or generate useful metabolites such as nitrates or tetrathionate ^22,30,33,35,59^. Indeed, the representative facultative anaerobic system studied here mediates rapid bacterial chemoattraction to O_2_ sources as well as low concentrations of ROS that can confer a metabolic advantage. As opposed to responding only to specific ROS, CZBs may offer bacteria a generalized mechanism of redox sensing by exploiting the high sensitivity of Zn^2+^ solubility, using the concentration of soluble Zn²⁺ as a proxy for environmental redox potential ^17,22,74,75^.

### A proposed model of sensing by CZB protein domains

We propose that our data are consistent with a model in which CZB domains mediate redox taxis through a Zn²⁺-dependent sensory mechanism in which the bioavailable Zn²⁺ pool serves as a secondary messenger for the redox state of the local environment (Fig. 4F). Importantly, this mechanism does not rely on Zn²⁺ undergoing oxidation or reduction. Rather, environmental redox chemistry alters the speciation and bioavailability of zinc by shifting the equilibrium between soluble Zn²⁺ and insoluble zinc-containing precipitates, and McpA senses the resulting changes in the soluble Zn²⁺ pool.

This work addresses a longstanding uncertainty over whether CZBs mediate attraction or repulsion to ROS, and whether these responses differ among oxidants. By conducting titrations of HOCl across several orders of magnitude, we in fact find a bimodal response in that extremely high source concentrations promote chemorepulsion but lower concentrations within physiologically relevant regimes promote chemoattraction across HOCl, H_2_O_2_, and ROOH (Fig. 1)^41^. ROS chemoattraction also occurs at concentrations below those that cause bactericidal effects, thereby offering a possible explanation for the apparent paradox of attraction to bactericidal oxidants (Fig. 3A-B). Further, these nanomolar-to-micromolar ROS concentrations elicit clear changes in swimming behavior consistent with chemoattraction responses in the form of reduced tumbling rates and increased migration (Fig. 2).

While swimming assays in homogenous effector baths have frequently been used as proxies for chemotaxis responses, particularly for bacteria in which other types of chemotaxis assays are technically challenging, we were conscious of the limitations in interpreting these behaviors without orthogonal experiments that directly visualize bacterial localization to a point source of the effector. In assessing the function of McpA in redox taxis, the swimming data paired with CIRA shows clearly how the distribution of the bacterial population changes over time in response to the treatment source. Further, by employing automated tracking, the swimming analyses are based upon assessment of thousands of tracks pooled from different experiments such that we have greater confidence in our measurements of how ROS, Zn^2+^ levels, and O_2_ impact swimming, and how this relates to bacterial localization. In this study, the two assays yielded mostly concordant conclusions: CIRA resolved the spatial and temporal redistribution of the population relative to a defined source, including repulsion and adaptation at high concentrations, whereas the swimming assays provided single-cell migration and tumbling metrics for the receptor mutants at defined bath concentrations. Our finding that McpA directs attracted to diverse oxidants may also help to explain our previous observation that depletion of host-derived HOCl had no effect on McpA-mediated fitness during colitis, as McpA responds not only to HOCl but also to hydroperoxides and elevated O₂, such that elimination of a single attractant signal would be insufficient to abolish McpA signaling ^32,83^.

If McpA senses exogenous redox potential through changes in the soluble Zn²⁺ pool, and also the zinc-binding Cys thiolate can be directly oxidized by high levels of HOCl, for such host-associated pathogens that elicit strong inflammatory responses like *Salmonella* that encounter high concentrations of neutrophil-derived HOCl, both mechanisms could be at play (Fig. 4F) ^33,34,39,40^. Based on earlier work we would expect direct cysteine oxidation to approximate the behavior of the C272A mutant, where the cysteine disengages the Zn^2+^ ligand, lowering the affinity of the zinc-binding site and promoting release ^17,20,22^. This may act as an additional regulatory mechanism for sensory tuning for the subset of CZB-possessing pathogens for which host-derived HOCl might serve as an important cue during pathogenesis (Fig. 4F).

More broadly, and likely of greatest relevance to the diverse bacteria that possess CZB proteins, the defining chemoselectivity of CZBs may be their ability to distinguish soluble Zn^2+^ from insoluble zinc species. Using simple Nernst-based estimates as a heuristic framework, we compared the theoretical magnitude of effective redox perturbations detected by the CZB Zn^2+^-dependent mechanism with those of more canonical cysteine-, flavin-, and heme-based redox sensors. Reported operating ranges for several redox-responsive systems are available, including ArcB (∼20-50 mV) ^85^, OxyR (∼30 mV) ^86^, and Yap1/Orp1 (∼30 mV) ^87^. If the lower limit of detection is approximated from the early timepoints of the aerotaxis experiments, in which McpA-mediated attraction occurred across a 0.06–0.5 nM gradient of solubilized O₂, the corresponding theoretical redox perturbation would be on the order of only 8 mV for the O₂/H₂O couple ^88^, suggesting, speculatively, this CZB system to be one of the most sensitive O_2_ sensors so far described.

## Materials and Methods

### Preparation of motile Salmonella enterica serovar Typhimurium

Bacterial strains and plasmids used in this study are listed in Table S1. To prepare cells for motility and chemotaxis assays, bacteria were cultured overnight in 2 ml of tryptone broth (TB) and, for antibiotic selection, 50 µg/ml ampicillin (TB + Amp), at 30 °C. 50 µl of overnight culture was used to inoculate 25 ml of fresh TB + Amp the next day, followed by shaking for 3 hours to reach approximately A_600_ of 0.5. Bacterial cultures were centrifuged at 1,500 g for 20 minutes and resuspended into chemotaxis buffer (CB) containing 10 mM potassium phosphate and 10 mM sodium lactate, and titrated to be pH 7. Cultures were then diluted to approximately A_600_ = 0.2 and rocked for 30-60 minutes before experimentation. For hypoxic CIRA experiments, the same protocol was used except that culturation was performed in an AnaeroJar 2.5L Jar (Oxoid, Basingstoke, UK) and a ThermoFisher CampyGen microaerophilic 2.5L Sachet (Waltham, Massachusetts, USA) with shaking at 37 °C.

### In vitro growth with ROS challenge

Wild-type *Salmonella* Typhimurium IR715 was incubated overnight in 25 ml of Luria-Bertani (LB) + Amp. The following day, cultures were pelleted by centrifugation at 2000 g for 20 minutes and resuspended in minimal media (MM) containing 47 mM Na_2_HPO_4_, 22 mM KH_2_PO_4_, 8 mM NaCl, 2mM MgSO_4_, 0.4% glucose (w/v) 11.35 mM (NH_4_)_2_SO_4_, and 100 μM CaCl_2_. 5 µl of the culture, adjusted to A_600_ = 2, was then used to inoculate fresh solutions of MM and additives of ROS (HOCl, H_2_O_2_, ROOH) in phosphate buffered saline (PBS), for a final volume of 200 µl per well in a 96-well microtiter plate. A Clariostar Plus plate reader (BMG Lab Tech, Ortenberg, DE) was used to monitor cell growth while shaking at 37° via A_600_ readings every 5 minutes for a period of 24 hours. Hypoxic growth experiments were conducted similarly under control of an atmospheric pressure unit (BMG Lab Tech, Ortenberg, DE) maintaining 10% CO_2_ and 1% O_2_ for the duration of data collection.

### Chemosensory injection rig assay (CIRA)

The CIRA apparatus is described previously ^89^. Briefly, treatment solutions are injected through a Femtotip II glass microcapillary (Eppendorf, Hamburg, DE) controlled with a MP-285 micromanipulator (Sutter, Novato, CA) into a 50 µl pond of motile bacteria on a 10-well slide (MP Biomedicals, Solon, OH). To generate a microgradient of effector, a constant flow of approximately 300 fL/min from the microcapillary was induced by applying compensation pressure (P_c_) of 35 hPa. Treatment solutions were made fresh each day, diluted into CB, and adjusted to pH 7. Bacterial responses were imaged with an inverted Nikon Ti2 Eclipse microscope (Tokyo, JP) with multichannel fluorescence imaging capabilities. For experiments done under hypoxic conditions, the unit was housed within a Coy Lab chamber (Grass Lake, Michigan, USA), at 1% O_2_, 10% CO_2_ and 37 °C.

### CIRA microgradient modeling

The CIRA microgradient was modeled according to the 3-dimension diffusion equation, as described previously ^44^. Diffusion coefficients for effectors were 6.5 x 10^-6^ cm^2^/sec for HOCl and H_2_O_2_, and 4.5 x 10^-6^ cm^2^/sec for cumene hydroperoxide, based on prior work ^90–92^.

### Quantification of CIRA data

Videos of chemotactic responses were quantified as described previously ^44^. The number of cells in each frame was calculated by determining a fluorescence intensity ratio per cell for frames pretreatment and calculating fluorescence intensity over time using the ‘plot profile’ function of ImageJ. Cell numbers were normalized to a baseline of ‘100%’ at the start of treatment (shown as time 0). Distribution of the bacterial population was quantified through use of the ‘radial profile’ ImageJ plugin. Radial distribution data were normalized by setting the field of view periphery as the baseline of ‘onefold’, which we defined as 280 µm distance from the source. For experiments with non-fluorescent cells, equivalent procedures were performed using phase-contrast data and enumeration of cells over time using a MATLAB-based tracking software, as described previously^53^. For hypoxic CIRA data, a photobleach correction was applied to all videos using the “Bleach Correction” plugin in ImageJ ^93^.

### Bacterial swimming assay

Bacteria were cultured as described above and back-diluted to A_600_ = 0.025 in a solution of CB or effector diluted into CB. This density allows for high-quality tracks of individual bacteria to be obtained with minimal overlapping of swimming trajectories. For each experiment, a fresh pond of 50 µl of motile bacteria was mounted on a 10-well slide (MP Biomedicals, Solon, OH). Bacteria were imaged over a 30 s period. Bacteria were then tracked using TrackingGUI ^53^, with bpfiltsize set to 20, the threshold option set to 2 (STD of thresholds), and the default threshold set at 4.0. Tracks were culled on the basis of the STD of position = 2, and the track length = the frame rate of the video (typically 25 fps), thereby selecting only motile cells and those that were monitored for at least one full second. Linear migration for the first second of each track was determined by:

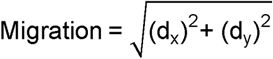

Where d_x_ and d_s_ are the differences in x and y pixel coordinates between the first and last valid frame within the first one-second window of each track. Average speed for the first second of each track was calculated as:

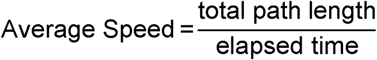

Where the total path length is every frame-to-frame Euclidean step distance within the first second of each track, and elapsed time is the actual elapsed time of those frames.

Finally, tumble detection was performed based on the approach described in Johnson et al. in 2024 ^94^. Briefly, instantaneous translational speed and angular speed were calculated from frame-to-frame positional differences and smoothed using a Savitzky-Golay filter of polynomial order 2 with a window automatically scaled to approximate 200 ms. Tumble candidates were identified as local minima in the smoothed speed trace. Each candidate was retained only if the difference between the adjacent peak speed and the minimum speed was at least 70% of the minimum speed itself. Candidates were then subject to two additional confirmation criteria. First, angular speed at the candidate frame (±1 frame) was required to exceed the track mean angular speed by at least 20%, consistent with the direction change expected during a genuine tumble event ^94^. Second, the speed at the minimum had to exceed 3.0 µm/s, to exclude artifactual tumbles arising from bacteria that stopped due to contact with a surface or exit from the focal plane. Tumble frames were delineated as those falling between the flanking local maxima whose smoothed speed remained within 30% of the speed range between the minimum and the surrounding peaks. Tumble frequency was then calculated for each bacterial swimming track as the total number of confirmed tumbles divided by total track duration in seconds.

### Free Zn^2+^ Assay

1 mM solutions of ZnSO_4_ were prepared under both hypoxic and normoxic conditions +/-the addition of ROS. After 3 hours of incubation, insoluble zinc species precipitated within the tube and were removed by centrifugation. The supernatant was subsequently removed and assayed with Zinpyr-1 (Santa Cruz Biotechnology, Dallas, TX, USA), a fluorescent probe that specifically binds Zn^2+^ ^95,96^. From there, the fluorescence of the solutions from 500-600 nm was determined based on a standard curve to determine the concentration of Zn^2+^ in each solution.

### Statistical Analysis

Data from replicate experiments were averaged and interpreted on the basis of their mean and standard error of the mean. Where relevant, p-values were calculated using Kruskal-Wallis test followed by Dunn’s multiple comparison tests or by unpaired t-tests, with significance determined at p<0.05.

## Data Availability

Because the raw video files reported in this study sometimes exceed 20 gigabytes each, totaling approximately 3 terabytes for all the videos in this study, it is not feasible to store all the video source files in a public repository. However, these data are available upon request from the corresponding author, Arden Baylink, at. Any of the data presented in our work will be shared with requesters promptly and without restriction. The video files are supplied in .avi or .nd2 format and can easily be viewed using the free ImageJ software, available here: https://imagej.net/ij/download.html. No other specialized software or code is needed to view or analyze the data.

## Acknowledgments

Funding for this work was provided by NIAID under award numbers 1K99AI148587 and 4R00AI148587-03, the EschLEAD program, Stanley Adler Fund, Autzen Foundation, and startup funding from Washington State University to AB. Bacterial strains were provided by Nikki Shariat (University of Georgia, Athens), Nkuchia Mikanatha and Pennsylvania NARMS and GenomeTrakr Programs, and Andreas Bäumler (University of California, Davis).

## Supplementary Materials

**Fig. S1.**
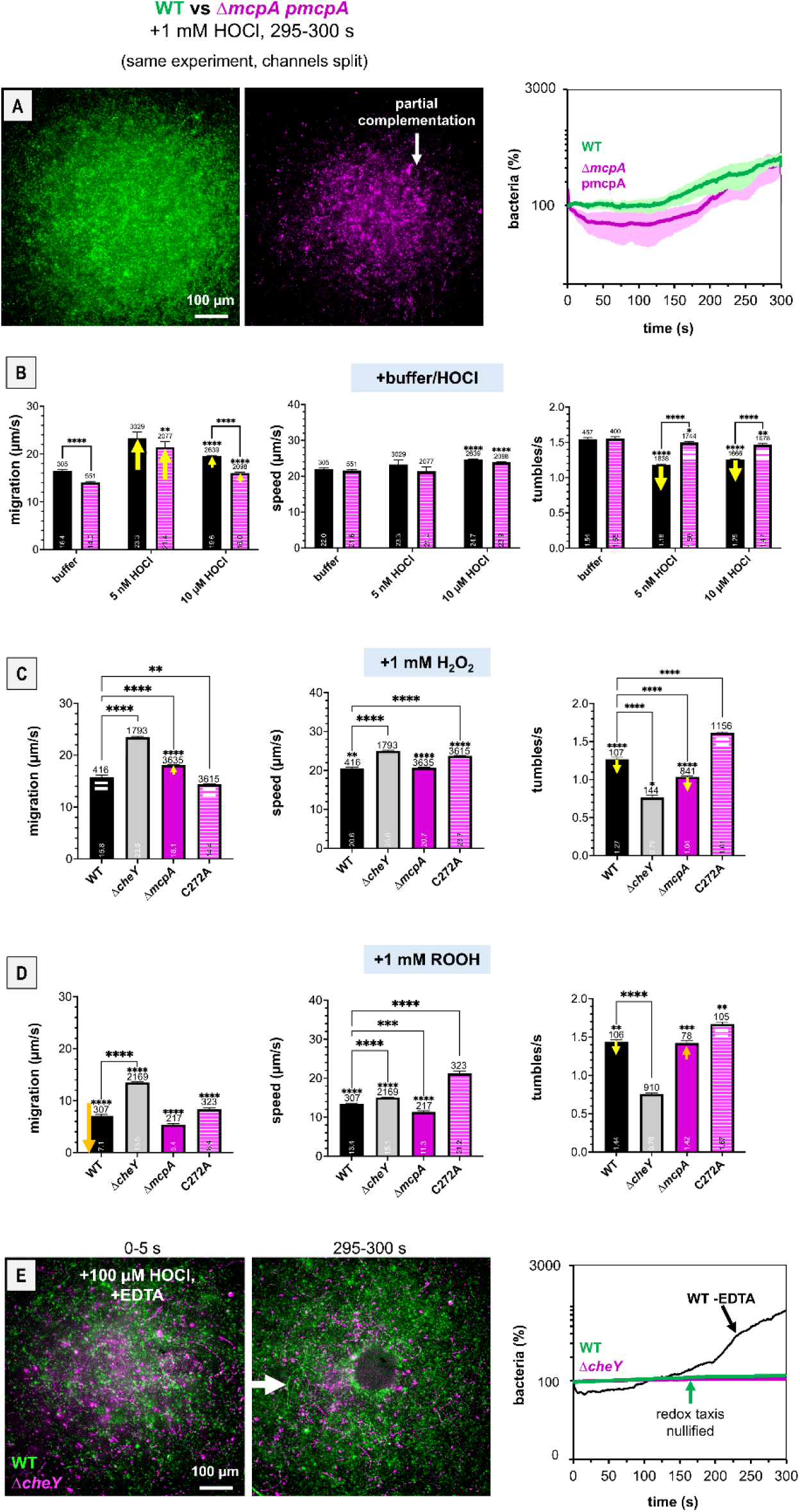
CIRA and swimming controls and supporting data A. Dual-channel imaging of chemotactic responses to 1 mM HOCl by WT (green) versus a Δ*mcpA* pmcpA complementation mutant (magenta); bacteria were co-cultured in the same experiment, but channels are separated for clarity. B-D. Swimming behavior in buffer or in the presence of ROS for WT IR715 (black), Δ*cheY* (gray), Δ*mcpA* (magenta), or C272A (magenta and white stripes). E. Dual-channel imaging of chemotactic responses to 1 mM HOCl by WT (green) and a Δ*cheY* mutant (magenta) cultured in chemotaxis buffer with 100 µM EDTA added, showing a null-response in this background. For reference, the WT response to HOCl without EDTA is shown in black. All CIRA data are means and error bars are standard error of the mean (SEM, n=3-5 Data means, error bars, and statistical calculations are as described in the main text.

## Supplementary Videos

**Supplementary Video 1:** Responses of WT *S.* Typhimurium IR715 and Δ*cheY* to 1 mM ROS at 10X speed. Experiments were conducted under normoxic conditions at 22°C. Full video length is 5 minutes. Viewable at: https://youtu.be/t56cjShQDZM

**Supplementary Video 2:** Responses of WT *S.* Typhimurium IR715 and Δ*mcpA* to 1 mM ROS at 10X speed. Experiments were conducted under normoxic conditions at 22°C. Full video length is 5 minutes. Viewable at: https://youtu.be/iOXzixrMu7M

**Supplementary Video 3:** Responses of WT *S.* Typhimurium IR715 and C272A to 1 mM ROS at 10X speed. Experiments were conducted under normoxic conditions at 22°C. Full video length is 5 minutes. Viewable at: https://youtu.be/ElE2J3BfM2s

**Supplementary Video 4:** Responses of WT *S.* Typhimurium and C272A to zinc-altered buffers (Chelex = zinc-chelated, Chelex +1 mM ZnSO_4_ = zinc-chelated with reintroduced Zn^2+^, +1 mM ZnO = zinc-chelated with added ZnO) at 10X speed. Experiments were conducted under normoxic conditions at 22°C. Full video length is 5 minutes. Viewable at: https://youtu.be/gfEmb01Rg2s

**Supplementary Video 5:** Responses of WT *S.* Typhimurium IR715 to injected normoxically equilibrated chemotaxis buffer shown at 7.5X speed. Bacteria were cultured prior to experiments under hypoxic (1% O_2_) conditions at 37 °C. Experiments were conducted under hypoxic conditions at 37°C. Full video length is 5 minutes. Viewable at: https://www.youtube.com/shorts/_zgOiq_G3fo

---

**Table S1.**
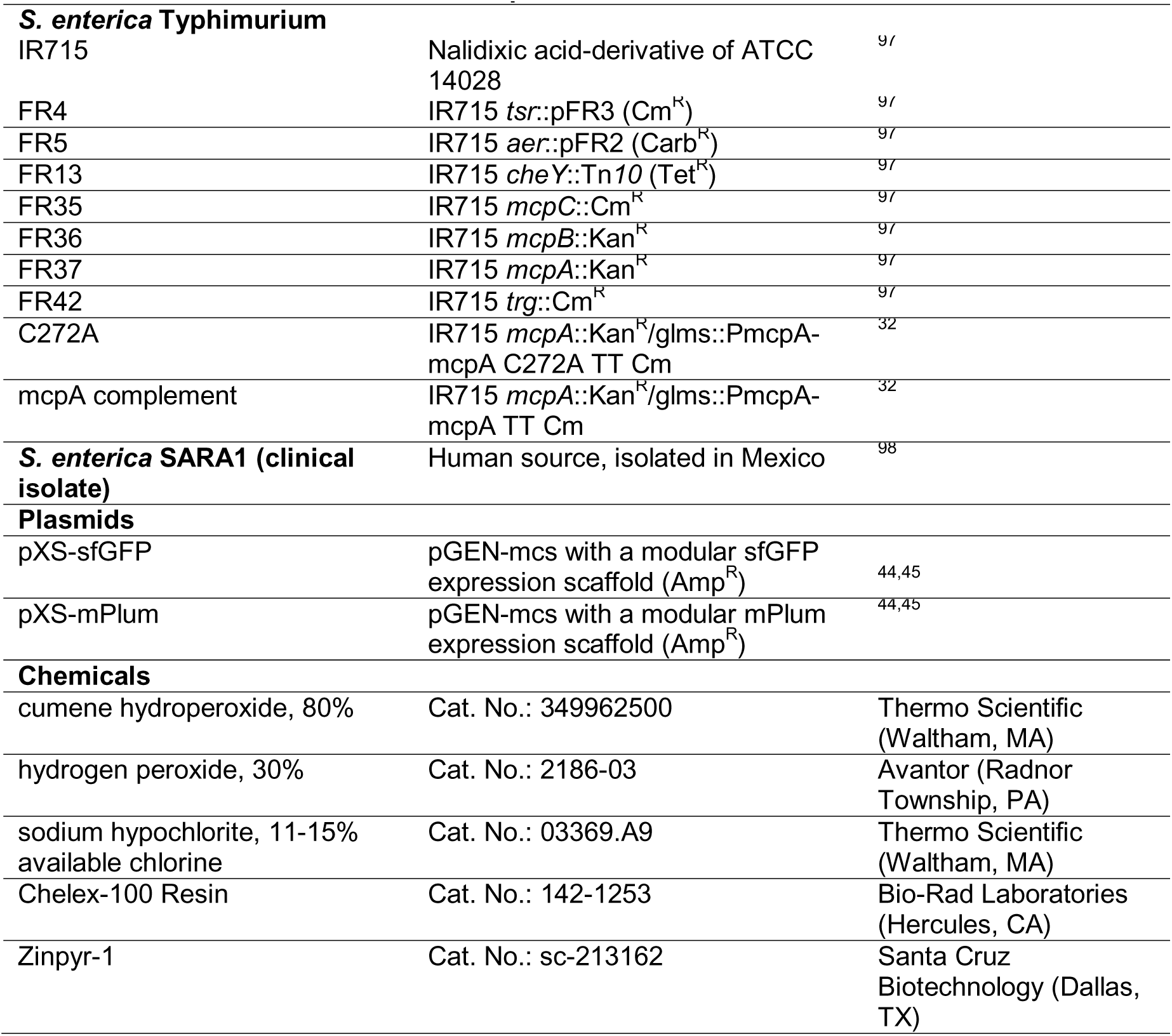
Bacterial strains, plasmids and chemicals used in this study.

## References

1. Olejarz, J., Iwasa, Y., Knoll, A.H., and Nowak, M.A. (2021). The Great Oxygenation Event as a consequence of ecological dynamics modulated by planetary change. Nat Commun 12, 3985.

2. Lyons, T.W., Reinhard, C.T., and Planavsky, N.J. (2014). The rise of oxygen in Earth’s early ocean and atmosphere. Nature 506, 307–315.

3. Checa, J., and Aran, J.M. (2020). Reactive Oxygen Species: Drivers of Physiological and Pathological Processes. J Inflamm Res 13, 1057–1073.

4. Epstein, F.H., and Weiss, S.J. (1989). Tissue Destruction by Neutrophils. N Engl J Med 320, 365–376.

5. Degrossoli, A. (2018). Neutrophil-generated HOCl leads to non-specific thiol oxidation in phagocytized bacteria. eLife 7, e32288.

6. Sies, H. (2022). Defining roles of specific reactive oxygen species (ROS) in cell biology and physiology. Nat Rev Mol Cell Biol 23, 499–515.

7. Perkins, A., Nelson, K.J., Parsonage, D., Poole, L.B., and Karplus, P.A. (2015). Peroxiredoxins: guardians against oxidative stress and modulators of peroxide signaling. Trends in Biochemical Sciences 40, 435–445. 10.1016/j.tibs.2015.05.001.

8. Sies, H., Mailloux, R.J., and Jakob, U. (2024). Fundamentals of redox regulation in biology. Nat Rev Mol Cell Biol 25, 701–719.

9. Sies, H., Belousov, V.V., Chandel, N.S., Davies, M.J., Jones, D.P., Mann, G.E., Murphy, M.P., Yamamoto, M., and Winterbourn, C. (2022). Defining roles of specific reactive oxygen species (ROS) in cell biology and physiology. Nat Rev Mol Cell Biol 23, 499–515. 10.1038/s41580-022-00456-z.

10. Murphy, M.P., Bayir, H., Belousov, V., Chang, C.J., Davies, K.J.A., Davies, M.J., Dick, T.P., Finkel, T., Forman, H.J., Janssen-Heininger, Y., et al. (2022). Guidelines for measuring reactive oxygen species and oxidative damage in cells and in vivo. Nat Metab 4, 651–662. 10.1038/s42255-022-00591-z.

11. Bibikov, S.I., Biran, R., Rudd, K.E., and Parkinson, J.S. (1997). A signal transducer for aerotaxis in Escherichia coli. J Bacteriol 179, 4075–4079. 10.1128/jb.179.12.4075-4079.1997.

12. Green, J., and Paget, M.S. (2004). Bacterial redox sensors. Nat Rev Microbiol 2, 954–966. 10.1038/nrmicro1022.

13. Kiley, P.J., and Beinert, H. (1998). Oxygen sensing by the global regulator, FNR: the role of the iron-sulfur cluster. FEMS Microbiol Rev 22, 341–352. 10.1111/j.1574-6976.1998.tb00375.x.

14. Malpica, R., Franco, B., Rodriguez, C., Kwon, O., and Georgellis, D. (2004). Identification of a quinone-sensitive redox switch in the ArcB sensor kinase. Proc Natl Acad Sci U S A 101, 13318–13323. 10.1073/pnas.0403064101.

15. Zheng, M., Aslund, F., and Storz, G. (1998). Activation of the OxyR transcription factor by reversible disulfide bond formation. Science 279, 1718–1721. 10.1126/science.279.5357.1718.

16. Ding, H., Hidalgo, E., and Demple, B. (1996). The redox state of the [2Fe-2S] clusters in SoxR protein regulates its activity as a transcription factor. J Biol Chem 271, 33173–33175. 10.1074/jbc.271.52.33173.

17. Perkins, A., Tudorica, D.A., Teixeira, R.D., Schirmer, T., Zumwalt, L., Ogba, O.M., Cassidy, C.K., Stansfeld, P.J., and Guillemin, K. (2021). A Bacterial Inflammation Sensor Regulates c-di-GMP Signaling, Adhesion, and Biofilm Formation. mBio 12. 10.1128/mBio.00173-21.

18. Franco, K., DiIorio, M., Simpkin, A.J., Keegan, R.M., Kallio, K., Goers-Sweeney, E., Colbert, M., Remington, S.J., Cassidy, C.K., Kulczyk, A.W., et al. (2026). Structure and signaling mechanism of Helicobacter pylori transducer-like protein D. Preprint at bioRxiv, 10.64898/2026.01.16.699579 10.64898/2026.01.16.699579.

19. Draper, J., Karplus, K., and Ottemann, K.M. (2011). Identification of a Chemoreceptor Zinc-Binding Domain Common to Cytoplasmic Bacterial Chemoreceptors. Journal of Bacteriology 193, 4338–4345. 10.1128/jb.05140-11.

20. Zähringer, F., Lacanna, E., Jenal, U., Schirmer, T., and Boehm, A. (2013). Structure and Signaling Mechanism of a Zinc-Sensory Diguanylate Cyclase. Structure 21, 1149–1157. 10.1016/j.str.2013.04.026.

21. Franco, K., DiIorio, M., Simpkin, A.J., Keegan, R., Kallio, K., Goers-Sweeney, E., Colbert, M., Remington, S.J., Cassidy, C.K., Kulczyk, A., et al. (2026). Structure and signaling mechanism of Helicobacter pylori transducer-like protein D. Preprint at bioRxiv, 10.64898/2026.01.16.699579 10.64898/2026.01.16.699579.

22. Perkins, A., Tudorica, D.A., Amieva, M.R., Remington, S.J., and Guillemin, K. (2019). Helicobacter pylori senses bleach (HOCl) as a chemoattractant using a cytosolic chemoreceptor. PLOS Biology 17, e3000395. 10.1371/journal.pbio.3000395.

23. Zumwalt, L., Perkins, A., and Ogba, O.M. (2020). Mechanism and Chemoselectivity for HOCl-Mediated Oxidation of Zinc-Bound Thiolates. ChemPhysChem 21, 2384–2387. 10.1002/cphc.202000634.

24. Collins, K.D., Andermann, T.M., Draper, J., Sanders, L., Williams, S.M., Araghi, C., and Ottemann, K.M. (2016). The Helicobacter pylori CZB Cytoplasmic Chemoreceptor TlpD Forms an Autonomous Polar Chemotaxis Signaling Complex That Mediates a Tactic Response to Oxidative Stress. J Bacteriol 198, 1563–1575. 10.1128/JB.00071-16.

25. Outten, C.E., and O’Halloran, T.V. (2001). Femtomolar sensitivity of metalloregulatory proteins controlling zinc homeostasis. Science 292, 2488–2492. 10.1126/science.1060331.

26. Capdevila, D.A., Wang, J., and Giedroc, D.P. (2016). Bacterial Strategies to Maintain Zinc Metallostasis at the Host-Pathogen Interface*. Journal of Biological Chemistry 291, 20858– 20868. 10.1074/jbc.R116.742023.

27. Helmann, J.D. (2025). Metals in Motion: Understanding Labile Metal Pools in Bacteria. Biochemistry 64, 329–345. 10.1021/acs.biochem.4c00726.

28. Collins, K.D., Hu, S., Grasberger, H., Kao, J.Y., and Ottemann, K.M. (2018). Chemotaxis Allows Bacteria To Overcome Host-Generated Reactive Oxygen Species That Constrain Gland Colonization. Infection and Immunity 86, 10.1128/iai.00878-17. 10.1128/iai.00878-17.

29. Behrens, W., Schweinitzer, T., McMurry, J.L., Loewen, P.C., Buettner, F.F.R., Menz, S., and Josenhans, C. (2016). Localisation and protein-protein interactions of the Helicobacter pylori taxis sensor TlpD and their connection to metabolic functions. Sci Rep 6, 23582. 10.1038/srep23582.

30. Zhou, B., Szymanski, C.M., and Baylink, A. (2022). Bacterial chemotaxis in human diseases. Trends in Microbiology. 10.1016/j.tim.2022.10.007.

31. Lebrun, V., Ravanat, J.-L., Latour, J.-M., and Sénèque, O. (2016). Near diffusion-controlled reaction of a Zn(Cys)4 zinc finger with hypochlorous acid. Chem. Sci 7, 5508–5516.

32. Cooper, K.G., Kari, L., Chong, A., Tandon, N., Doran, K., Gomes Da Silva, L., Cockrell, D.C., Baylink, A., and Steele-Mortimer, O. HilD-regulated chemotaxis proteins contribute to Salmonella Typhimurium colonization in the gut. mBio 16, e00390–25. 10.1128/mbio.00390-25.

33. Rivera-Chávez, F., Winter, S.E., Lopez, C.A., Xavier, M.N., Winter, M.G., Nuccio, S.-P., Russell, J.M., Laughlin, R.C., Lawhon, S.D., Sterzenbach, T., et al. (2013). Salmonella Uses Energy Taxis to Benefit from Intestinal Inflammation. PLoS Pathogens 9, e1003267. 10.1371/journal.ppat.1003267.

34. Winter, S.E. (2010). Gut inflammation provides a respiratory electron acceptor for Salmonella. Nature 467, 426–429.

35. Rivera-Chávez, F. (2016). Energy Taxis toward Host-Derived Nitrate Supports a Salmonella Pathogenicity Island 1-Independent Mechanism of Invasion. mBio 7, 00960–16.

36. Rolig, A.S., Shanks, J., Carter, J.E., and Ottemann, K.M. (2012). Helicobacter pylori Requires TlpD-Driven Chemotaxis To Proliferate in the Antrum. Infection and Immunity 80, 3713–3720. 10.1128/iai.00407-12.

37. Behrens, W., Schweinitzer, T., Bal, J., Dorsch, M., Bleich, A., Kops, F., Brenneke, B., Didelot, X., Suerbaum, S., and Josenhans, C. (2013). Role of Energy Sensor TlpD of Helicobacter pylori in Gerbil Colonization and Genome Analyses after Adaptation in the Gerbil. Infection and Immunity 81, 3534–3551. 10.1128/iai.00750-13.

38. Gül, E. (2024). Salmonella T3SS-2 virulence enhances gut-luminal colonization by enabling chemotaxis-dependent exploitation of intestinal inflammation. Cell Rep 43, 113925.

39. Thiennimitr, P. (2011). Intestinal inflammation allows Salmonella to use ethanolamine to compete with the microbiota. Proc. Natl. Acad. Sci. U.S.A *108*, 17480–17485.

40. Winter, S.E., Lopez, C.A., and Bäumler, A.J. (2013). The dynamics of gut-associated microbial communities during inflammation. EMBO Reports 14, 319–327.

41. Winterbourn, C.C., Hampton, M.B., Livesey, J.H., and Kettle, A.J. (2006). Modeling the Reactions of Superoxide and Myeloperoxidase in the Neutrophil Phagosome: IMPLICATIONS FOR MICROBIAL KILLING*. Journal of Biological Chemistry 281, 39860– 39869.

42. Chao, M.S. (1968). The Diffusion Coefficients of Hypochlorite, Hypochlorous Acid, and Chlorine in Aqueous Media by Chronopotentiometry. J. Electrochem. Soc 115, 1172.

43. Huang, D., Han, M., Wang, J., and Jin, Y. (2002). Catalytic decomposition process of cumene hydroperoxide using sulfonic resins as catalyst. Chemical Engineering Journal 88, 215–223.

44. Glenn, S.J., Gentry-Lear, Z., Shavlik, M., Harms, M.J., Asaki, T.J., and Baylink, A. (2024). Bacterial vampirism mediated through taxis to serum. eLife 12, RP93178. 10.7554/eLife.93178.

45. Franco, K., Gentry-Lear, Z., Shavlik, M., Harms, M.J., and Baylink, A. (2025). Navigating contradictions in enteric chemotactic stimuli. eLife 14, RP106261. 10.7554/eLife.106261.

46. Howitt, M.R., Lee, J.Y., Lertsethtakarn, P., Vogelmann, R., Joubert, L.-M., Ottemann, K.M., and Amieva, M.R. (2011). ChePep Controls Helicobacter pylori Infection of the Gastric Glands and Chemotaxis in the Epsilonproteobacteria. mBio 2, e00098–11. 10.1128/mBio.00098-11.

47. Bi, S., and Lai, L. (2015). Bacterial chemoreceptors and chemoeffectors. Cellular and Molecular Life Sciences 72, 691–708. 10.1007/s00018-014-1770-5.

48. Berg, H.C., and Brown, D.A. (1972). Chemotaxis in Escherichia coli analysed by Three-dimensional Tracking. Nature 239, 500–504. 10.1038/239500a0.

49. Bi, S., and Sourjik, V. (2018). Stimulus sensing and signal processing in bacterial chemotaxis. Current Opinion in Microbiology 45, 22–29. 10.1016/j.mib.2018.02.002.

50. Grognot, M., and Taute, K.M. (2021). A multiscale 3D chemotaxis assay reveals bacterial navigation mechanisms. Commun Biol 4, 669. 10.1038/s42003-021-02190-2.

51. Hazelbauer, G.L., and Parkinson, J.S. (1977). Bacterial Chemotaxis. In Microbial Interactions, J. L. Reissig, ed. (Springer US), pp. 59–98. 10.1007/978-1-4615-9698-1_2.

52. Hazelbauer, G.L., Falke, J.J., and Parkinson, J.S. (2008). Bacterial chemoreceptors: high-performance signaling in networked arrays. Trends in Biochemical Sciences 33, 9–19. 10.1016/j.tibs.2007.09.014.

53. Parthasarathy, R. (2012). Rapid, accurate particle tracking by calculation of radial symmetry centers. Nat Methods 9, 724–726. 10.1038/nmeth.2071.

54. Bhattarai, N., Moore, J.P., Payne, S., and Harshey, R.M. (2026). Aer is a bidirectional redox sensor mediating negative chemotaxis to antibiotic-induced ROS in Escherichia coli. mBio 17, e00381–26. 10.1128/mbio.00381-26.

55. Greer-Phillips, S.E., Alexandre, G., Taylor, B.L., and Zhulin, I.B. (2003). Aer and Tsr guide Escherichia coli in spatial gradients of oxidizable substrates. Microbiology (Reading) 149, 2661–2667. 10.1099/mic.0.26304-0.

56. Rebbapragada, A., Johnson, M.S., Harding, G.P., Zuccarelli, A.J., Fletcher, H.M., Zhulin, I.B., and Taylor, B.L. (1997). The Aer protein and the serine chemoreceptor Tsr independently sense intracellular energy levels and transduce oxygen, redox, and energy signals for Escherichia coli behavior. Proceedings of the National Academy of Sciences 94, 10541– 10546. 10.1073/pnas.94.20.10541.

57. Zheng, L., Kelly, C.J., and Colgan, S.P. (2015). Physiologic hypoxia and oxygen homeostasis in the healthy intestine. A Review in the Theme: Cellular Responses to Hypoxia. Am J Physiol Cell Physiol 309, C350–C360. 10.1152/ajpcell.00191.2015.

58. Singhal, R., and Shah, Y.M. (2020). Oxygen battle in the gut: Hypoxia and hypoxia-inducible factors in metabolic and inflammatory responses in the intestine. J Biol Chem 295, 10493– 10505. 10.1074/jbc.REV120.011188.

59. Miller, B.M. (2020). Anaerobic respiration of NOX1-derived hydrogen peroxide licenses bacterial growth at the colonic surface. Cell Host Microbe 28, 789–797 5.

60. Khademian, M., and Imlay, J.A. (2017). Escherichia coli cytochrome c peroxidase is a respiratory oxidase that enables the use of hydrogen peroxide as a terminal electron acceptor. Proc Natl Acad Sci U S A 114, E6922–E6931. 10.1073/pnas.1701587114.

61. Wenegieme, T.-Y., Elased, D., McMichael, K.E., Rockwood, J., Hasrat, K., Ume, A.C., Marshall, A.G., Neikirk, K., Kirabo, A., Elased, K.M., et al. (2024). Strategies for inducing and validating zinc deficiency and zinc repletion. American Journal of Physiology-Heart and Circulatory Physiology 326, H1396–H1401. 10.1152/ajpheart.00025.2024.

62. Zinc Dioxide Nanoparticulates: A Hydrogen Peroxide Source at Moderate pH | Environmental Science & Technology https://pubs.acs.org/doi/full/10.1021/es4020629.

63. Jomova, K., Alomar, S.Y., Valko, R., Nepovimova, E., Kuca, K., and Valko, M. (2025). The role of redox-active iron, copper, manganese, and redox-inactive zinc in toxicity, oxidative stress, and human diseases. EXCLI J 24, 880–954. 10.17179/excli2025-8449.

64. Lee, S.R. (2018). Critical Role of Zinc as Either an Antioxidant or a Prooxidant in Cellular Systems. Oxidative Medicine and Cellular Longevity 2018, 9156285. 10.1155/2018/9156285.

65. Moezzi, A., Cortie, M., and McDonagh, A. (2011). Aqueous pathways for the formation of zinc oxide nanoparticles. Dalton Trans 40, 4871.

66. Oteiza, P.I. (2012). Zinc and the modulation of redox homeostasis. Free Radic Biol Med 53, 1748–1759. 10.1016/j.freeradbiomed.2012.08.568.

67. Alker, W., Schwerdtle, T., Schomburg, L., and Haase, H. (2019). A Zinpyr-1-based Fluorimetric Microassay for Free Zinc in Human Serum. Int J Mol Sci 20, 4006. 10.3390/ijms20164006.

68. Jurinak, J.J. (1976). The Hydrolysis of Cations. Soil Science Society of America Journal 40, vi–vi. 10.2136/sssaj1976.03615995004000030006x.

69. Krężel, A., and Maret, W. (2016). The biological inorganic chemistry of zinc ions. Archives of Biochemistry and Biophysics 611, 3–19. 10.1016/j.abb.2016.04.010.

70. Taylor, B.L., Zhulin, I.B., and Johnson, M.S. (1999). Aerotaxis and other energy-sensing behavior in bacteria. Annu Rev Microbiol 53, 103–128. 10.1146/annurev.micro.53.1.103.

71. PeerJ Bacterial aerotaxis: receptor diversity, signal transduction and ecological functions https://peerj.com/articles/21573/.

72. Alexandre, G. (2010). Coupling metabolism and chemotaxis-dependent behaviours by energy taxis receptors. Microbiology 156, 2283–2293. 10.1099/mic.0.039214-0.

73. Brunet, M., Amin, S.A., Bodachivskyi, I., Kuzhiumparambil, U., Seymour, J.R., and Raina, J.-B. (2025). An atlas of metabolites driving chemotaxis in prokaryotes. Nat Commun 16, 1242. 10.1038/s41467-025-56410-y.

74. Hübner, C., and Haase, H. (2021). Interactions of zinc-and redox-signaling pathways. Redox Biology 41, 101916. 10.1016/j.redox.2021.101916.

75. Magerand, R., Rey, P., Blanchard, L., and de Groot, A. (2021). Redox signaling through zinc activates the radiation response in Deinococcus bacteria. Sci Rep 11, 4528. 10.1038/s41598-021-84026-x.

76. Vázquez-Torres, A. (2012). Redox Active Thiol Sensors of Oxidative and Nitrosative Stress. Antioxidants & Redox Signaling 17, 1201–1214. 10.1089/ars.2012.4522.

77. Janda, I., Devedjiev, Y., Derewenda, U., Dauter, Z., Bielnicki, J., Cooper, D.R., Graf, P.C.F., Joachimiak, A., Jakob, U., and Derewenda, Z.S. (2004). The Crystal Structure of the Reduced, Zn2+-Bound Form of the *B. subtilis* Hsp33 Chaperone and Its Implications for the Activation Mechanism. Structure *12*, 1901–1907. 10.1016/j.str.2004.08.003.

78. Graumann, J., Lilie, H., Tang, X., Tucker, K.A., Hoffmann, J.H., Vijayalakshmi, J., Saper, M., Bardwell, J.C.A., and Jakob, U. (2001). Activation of the Redox-Regulated Molecular Chaperone Hsp33—A Two-Step Mechanism. Structure 9, 377–387. 10.1016/S0969-2126(01)00599-8.

79. Zhao, F., Ilbert, M., Varadan, R., Cremers, C.M., Hoyos, B., Acin-Perez, R., Vinogradov, V., Cowburn, D., Jakob, U., and Hammerling, U. (2011). Are Zinc-Finger Domains of Protein Kinase C Dynamic Structures That Unfold by Lipid or Redox Activation? Antioxidants & Redox Signaling 14, 757–766. 10.1089/ars.2010.3773.

80. Antelmann, H., and Helmann, J.D. (2011). Thiol-Based Redox Switches and Gene Regulation. Antioxidants & Redox Signaling 14, 1049–1063. 10.1089/ars.2010.3400.

81. Cremers, C.M., and Jakob, U. (2013). Oxidant Sensing by Reversible Disulfide Bond Formation*. Journal of Biological Chemistry 288, 26489–26496. 10.1074/jbc.R113.462929.

82. McCord, M.C., and Aizenman, E. (2014). The role of intracellular zinc release in aging, oxidative stress, and Alzheimer’s disease. Front. Aging Neurosci. 6. 10.3389/fnagi.2014.00077.

83. Cooper, K.G., Chong, A., Kari, L., Jeffrey, B., Starr, T., Martens, C., McClurg, M., Posada, V.R., Laughlin, R.C., Whitfield-Cargile, C., et al. (2021). Regulatory protein HilD stimulates Salmonella Typhimurium invasiveness by promoting smooth swimming via the methyl-accepting chemotaxis protein McpC. Nature Communications 12, 348. 10.1038/s41467-020-20558-6.

84. Winter, J., Ilbert, M., Graf, P.C.F., Ozcelik, D., and Jakob, U. (2008). Bleach activates a redox-regulated chaperone by oxidative protein unfolding. Cell 135, 691–701. 10.1016/j.cell.2008.09.024.

85. Bekker, M., Alexeeva, S., Laan, W., Sawers, G., Teixeira de Mattos, J., and Hellingwerf, K. (2010). The ArcBA Two-Component System of Escherichia coli Is Regulated by the Redox State of both the Ubiquinone and the Menaquinone Pool. J Bacteriol 192, 746–754. 10.1128/JB.01156-09.

86. Åslund, F., Zheng, M., Beckwith, J., and Storz, G. (1999). Regulation of the OxyR transcription factor by hydrogen peroxide and the cellular thiol—disulfide status. Proceedings of the National Academy of Sciences 96, 6161–6165. 10.1073/pnas.96.11.6161.

87. Mason, J.T., Kim, S.-K., Knaff, D.B., and Wood, M.J. (2006). Thermodynamic Basis for Redox Regulation of the Yap1 Signal Transduction Pathway. Biochemistry 45, 13409–13417. 10.1021/bi061136y.

88. Thauer, R.K., Jungermann, K., and Decker, K. (1977). Energy conservation in chemotrophic anaerobic bacteria. Bacteriol Rev 41, 100–180. 10.1128/br.41.1.100-180.1977.

90. Chao, M.S. (1968). The Diffusion Coefficients of Hypochlorite, Hypochlorous Acid, and Chlorine in Aqueous Media by Chronopotentiometry. J. Electrochem. Soc. 115, 1172. 10.1149/1.2410933.

91. Huang, D., Han, M., Wang, J., and Jin, Y. (2002). Catalytic decomposition process of cumene hydroperoxide using sulfonic resins as catalyst. Chemical Engineering Journal 88, 215–223. 10.1016/S1385-8947(02)00003-7.

92. McMurtrie, R.L., and Keyes, F.G. (1948). A Measurement of the Diffusion Coefficient of Hydrogen Peroxide Vapor into Air. J. Am. Chem. Soc. 70, 3755–3758. 10.1021/ja01191a063.

93. Miura, K. (2020). Bleach correction ImageJ plugin for compensating the photobleaching of time-lapse sequences. Preprint at F1000Research, 10.12688/f1000research.27171.1 10.12688/f1000research.27171.1.

94. Johnson, S., Freedman, B., and Tang, J.X. (2024). Run-and-tumble kinematics of Enterobacter Sp. SM3. Phys. Rev. E *109*, 064402. 10.1103/PhysRevE.109.064402.

95. Sinclair, S.A., Sherson, S.M., Jarvis, R., Camakaris, J., and Cobbett, C.S. (2007). The use of the zinc-fluorophore, Zinpyr-1, in the study of zinc homeostasis in Arabidopsis roots. New Phytologist *174*, 39–45. 10.1111/j.1469-8137.2007.02030.x.

96. Figueroa, J.A.L., Vignesh, K.S., Deepe, G.S., and Caruso, J. (2014). Selectivity and specificity of small molecule fluorescent dyes/probes used for the detection of Zn2+ and Ca2+ in cells. Metallomics 6, 301–315. 10.1039/c3mt00283g.

97. Rivera-Chávez, F., Winter, S.E., Lopez, C.A., Xavier, M.N., Winter, M.G., Nuccio, S.-P., Russell, J.M., Laughlin, R.C., Lawhon, S.D., Sterzenbach, T., et al. (2013). Salmonella uses energy taxis to benefit from intestinal inflammation. PLoS Pathog 9, e1003267. 10.1371/journal.ppat.1003267.

98. Beltran, P., Plock, S.A., Smith, N.H., Whittam, T.S., Old, D.C., and Selander, R.K. (1991). Reference collection of strains of the Salmonella typhimurium complex from natural populations. Journal of General Microbiology 137, 601–606. 10.1099/00221287-137-3-601.

